# Jerusalem artichoke extracts regulate the gene expression of key enzymes involved in fatty acid biosynthesis

**DOI:** 10.1101/2024.09.30.615848

**Authors:** Heeyoun Bunch, Young-Ran Yoon, Stuart K. Calderwood

**Affiliations:** Department of Applied Biosciences, Kyungpook National University, Daegu 41566, Republic of Korea; School of Medicine, Kyungpook National University, Daegu 41944, Republic of Korea; Department of Radiation Oncology, Beth Israel Deaconess Medical Center, Harvard Medical School, Boston 02115, USA

**Keywords:** Gene regulation, Jerusalem artichoke, functional food, fatty acid biosynthesis, fatty acid synthase, acetyl-coA carboxylase, topoisomerase II

## Abstract

Jerusalem artichoke (JA) is a traditional remedy for alleviating symptoms of diabetes. In fact, the suppressive effects of JA on blood sugar have been reported in multiple studies since 1934. Recent studies have indicated that type II diabetes is often caused by insulin resistance rather than insulin reduction and that increased blood and interstitial fatty acid levels contribute to insulin resistance and the development of diabetes. However, whether JA affects lipogenesis has not been studied. Here, we elucidated the effects of JA on the expression of two key enzymes involved in fatty acid biosynthesis, fatty acid synthase (*FASN*) and acetyl-coA carboxylase (*ACACA*), using three immortalized human bone marrow, colon, and liver cell lines. Caffeine and ICRF193, a catalytic inhibitor of topoisomerase II (TOP2), were included as positive controls, and JA was extracted into water- or dimethyl sulfoxide-soluble components, termed H-JA and D-JA. D-JA significantly reduced the expression of *FASN* and *ACACA* at the mRNA and protein levels. D-JA-treated cells exhibited altered TOP2 levels and *FASN*/*ACACA* expression appeared to be controlled by TOP2 activity and levels. Taken together, our study revealed a novel effect of JA extracts on inhibiting the expression of the key enzymes involved in the fatty acid synthesis and suggested the potential of JA as a natural medicinal agent to control lipogenesis in humans.

**Highlights:**

- Jerusalem artichoke extracts reduce the expression of *FASN* and *ACACA* genes at the mRNA and protein levels
- Topoisomerase II regulates *FASN* and *ACACA* gene expression
- Jerusalem artichoke extracts and plasma glucose concentrations regulate cellular topoisomerase II protein expression

## INTRODUCTION

In modern societies, the incidence of metabolic syndrome and associate diseases, such as diabetes mellitus and stroke, has been increasing^1-3^. Metabolic syndrome is characterized by high blood sugar, high blood pressure, and increased body fat around the waist. Increased cholesterol and triglyceride levels also serve as the signature of the syndrome^4^. The exact causes of metabolic syndrome are not completely understood; however, poor diet, genetic preposition, and sedative lifestyle may contribute to the development of the syndrome^5,6^.

Humans require homeostasis of blood glucose levels at approximately 80–100 mg/dL (4–6 mM)^7^, which is crucial for maintaining healthy organs. Either higher or lower levels of glucose trigger the secretion of pancreatic insulin or glucagon hormones and turn on or off accordant catabolic and anabolic pathways of the body, as organismal efforts to restore the proper level of blood glucose^7,8^. After the digestion of food, monosaccharide glucose is circulated in the blood at increased levels and is taken up and actively metabolized by cells throughout the body. Pancreatic insulin secreted by high blood sugar levels ensures facilitated glucose transport into the cell, which is mediated by glucose transporters^9^. If this process is compromised and so blood glucose levels are not restored to the normal range, the condition is called hyperglycemia or diabetes. In type II diabetes, often referred to as adult diabetes, insulin resistance or abnormal secretion/production from pancreatic β cells results in hyperglycemia^10^. Insulin resistance is caused by impaired insulin receptor sensitivity to insulin. The exact causes of insulin resistance have been incompletely understood. However, excess blood glucose, obesity-mediated inflammation, and bodily inactivity have been reported to cause insulin resistance^11-13^. In addition, increased plasma triglyceride levels have been linked to insulin resistance^14^.

Jerusalem artichoke (*Helianthus tuberosus*; JA, hereafter) is a species of sunflower and a root vegetable. It originated in North America and is now cultivated in North and South America, Europe, Australia, and Asia. Because the tuber of JA contains little starch and a high content of inulin, a polymer of fructose that cannot be digested and absorbed in the human intestine to spike up blood glucose levels, it has been used to treat diabetes^15,16^. Inulin is also known to be a prebiotic that promotes the growth of beneficial gastrointestinal bacteria. For example, recent studies reported that consumption of grated JA before meals reduced plasma glucose^17^ and that inulin supplementation reduced hyperglycemia by improving intestinal microflora in a mouse model^16^. An interesting study also found that JA can prevent the onset of diabetes and non-alcoholic fatty liver disease in rats^18^.

Although the effects of JA on the prevention and alleviation of diabetes and the effects of plasma triglycerides on insulin resistance have been reported by multiple studies^14-16,18^, the effects of JA on fatty acid metabolism have not been studied. Altered or impaired fatty acid metabolism increases the levels of triglycerides, a covalent complex of a glycerol and three fatty acids, in the human body^19^.

Excess glucose/fructose consumption can stimulate the biosynthesis of fatty acids and triglycerides^20,21^. Fatty acid biosynthesis involves two key enzymes: acetyl-coA acetylase (*ACACA* or *ACC1*) and fatty acid synthase (*FASN* or *FAS*)^22^. ACACA converts acetyl-coA from glucose or amino acids into malonyl-coA, an important intermediate in fatty acid synthesis^22^. FASN regulates palmitoylation, thereby generating fatty acids^22^. In this study, we attempted to evaluate the potential role of JA in regulating *FASN* and *ACACA* gene expression, which can significantly affect fatty acid and triglyceride levels in humans.

Natural chemicals found in foods exert potent effects on gene regulation in a similar manner as extracellular signals. Compared with synthetic chemicals, food chemicals have many benefits as they are safer, easily incorporated into routine diets, and less addictive. In addition, given that these chemicals affect gene expression, their long-term effects can be expected to prevent and improve disease symptoms.

This study found that dimethyl sulfoxide-soluble JA (D-JA), but not water-soluble JA (H-JA), significantly reduces *FASN* and *ACACA* expression in three human cell lines. JA extracts are innocuous because both H-JA and D-JA do not exhibit cytotoxicity in our experimental concentration ranges. Despite some tissue-specific responses to D-JA regarding *FASN* expression, the overall effect of D-JA in suppressing *FASN* expression is comparable to that of the topoisomerase II (TOP2) catalytic inhibitor ICRF193^23^. We included this chemical for a previous report that one of the TOP2 isomers TOP2B is important for *FASN* transcription^24^. D-JA also effectively represses *ACACA* expression at both the mRNA and protein levels. In addition, we found that TOP2B levels fluctuate with glucose concentrations and determine the expression of *FASN* and *ACACA* genes. These findings suggest the fundamental gene regulatory and medicinal potential of JA for preventing and treating insulin resistance and abnormal fatty acid metabolism.

## MATERIALS & METHODS

### Cell culture and conditions

SH-SY5Y, HCT116, and HepG2 cells (American Type Culture Collection, USA) were grown in high-glucose DMEM, (Cytiva), MEM alpha modification (Hyclone), and DMEM (Corning) supplemented with 10% fetal bovine serum (FBS, Gibco) and 1% penicillin/streptomycin (P/S, Thermo Fisher) solution in a 5% CO_2_ incubator at 37°C. The cells were grown to 70%–80% confluence in a 10 cm dish before being split into 6 or 96 well plates. H-JA (H_2_O, control) and D-JA (DMSO, control) were applied at the indicated concentrations for 24–72 h. Caffeine (Sigma, C0750) and ICRF193 (Sigma, I4659) were treated at the indicated final concentrations along with H_2_O and DMSO as controls.

### Cytotoxicity test

SH-SY5Y, HCT116, and HepG2 cells were grown to approximately 50%–60% confluence in a 96-well plate containing complete media, the media supplemented with 10% FBS and 1% P/S. The media were exchanged with fresh one, including H-JA or D-JA in H_2_O or DMSO, respectively, 1% of the total media volume. JA-treated cells were incubated for 48–72 h before supplementing each well with 10% water-soluble tetrazolium salt (WST, DoGenBio, South Korea) according to the manufacturer’s instructions. The reaction was developed for 1–2 h and the absorbance at 450 nm was measured using a microplate reader (Tecan, Switzerland).

### Immunoblotting

Cells were grown to approximately 70%–80% confluence in 6-well plates. The controls and JA-treated cells were incubated for 72 h, washed with cold PBS, and then collected with RIPA buffer (Cell Signaling Technology). The protein concentrations of the cell lysates were measured using the Bradford assay (Bio-Rad). From the measured protein concentrations, equal amounts of proteins were loaded onto an SDS-polyacrylamide gel. The separated proteins were transferred to a nitrocellulose membrane (GE Healthcare). Primary antibodies were used for probing β-ACTIN (MA5-15739, Invitrogen), FASN (ab128856, Abcam), ACACA (#3662, Cell Signaling Technology), TOP2B (A300-949A, Bethyl Laboratories; sc-25330, Santa Cruz Biotechnology), DNA-PK (ab168854, Abcam), and α-tubulin (sc-8035, SantaCruz Technology). Each antibody was diluted in 5% skim milk (Bio-Rad) solution. After the primary antibody incubation, the membranes were rinsed twice and washed four times, each for 10 min, with PBST solution [137 mM NaCl, 2.7 mM KCl, 4.3 mM Na_2_HPO_4_, 1.5 mM KH_2_PO_4,_ 0.1% (v/v) Tween-20 and pH 7.4]. HRP-conjugated rabbit (#7074, Cell Signaling Technology) or mouse (#7076, Cell Signaling Technology) secondary antibodies, were diluted in 5% skim milk and incubated with the membranes. After incubation, the membranes were rinsed twice and washed four times with PBST solution, each for 10 min. Western Blotting Luminol Reagent (sc-2048, SantaCruz Technology) or SuperSignal West Atto Ultimate Sensitivity Substrate (A38554, Thermo Fisher) was used to develop signals.

### Quantitative real-time PCR (qRT–PCR)

cDNAs were constructed from 600 ng of the collected RNA via reverse transcription using ReverTra Ace qPCR RT Master Mix (Toyobo). Equal amounts of the resultant cDNAs were analyzed via real-time qPCR using Applied Biosystems PowerUp SYBR Green Master Mix and according to the manufacturer’s instructions (Applied Biosystems). The StepOnePlus Real-Time PCR System was used (Thermo Fisher Scientific). The thermal cycle used was 10 min for pre-denaturation followed by 45 cycles of 95°C for 15 s, 55°C for 15 s, and 72°C for 45 s. The primers used in the experiments were purchased from Integrated DNA technology and are summarized in **Supplementary Table 1**.

### Statistical analysis

The standard deviation was calculated and used to generate error bars. Student’s t-test (unpaired, one-way) was conducted to determine statistical significance. The graphs were generated using Prism 9 software (GraphPad, Inc.).

## RESULTS

### Generation of water- and DMSO-soluble JA extracts

As described in the materials and methods section in details, JA was sliced, freeze-dried, and then powdered. H-JA and D-JA were generated by extracting JA using water and DMSO, respectively, so that the polar and nonpolar compounds could be enriched by these distinctive solvents. Insoluble debris was removed via centrifugation and filtration (**Fig. 1A**). The chemical compositions of H-JA and D-JA were compared by gas chromatography-mass spectrometry (GC-MS) analysis. In total, 96 and 55 compounds were identified for H-JA and D-JA, respectively, including 12 common compounds (**Fig. 1B**; **Supplementary Data 1**). The unique compounds found in H-JA included hexanal and benzaldehyde, and in D-JA, hexadecane, 6,11-dipentyl-, and benz[f]isoquinoline, 5,6-dihydro-2,4,- dimethyl- (**Supplementary Data 1**). These data indicate that our extraction method effectively generated two chemically distinctive JA samples, H-JA and D-JA.

**Fig. 1.**
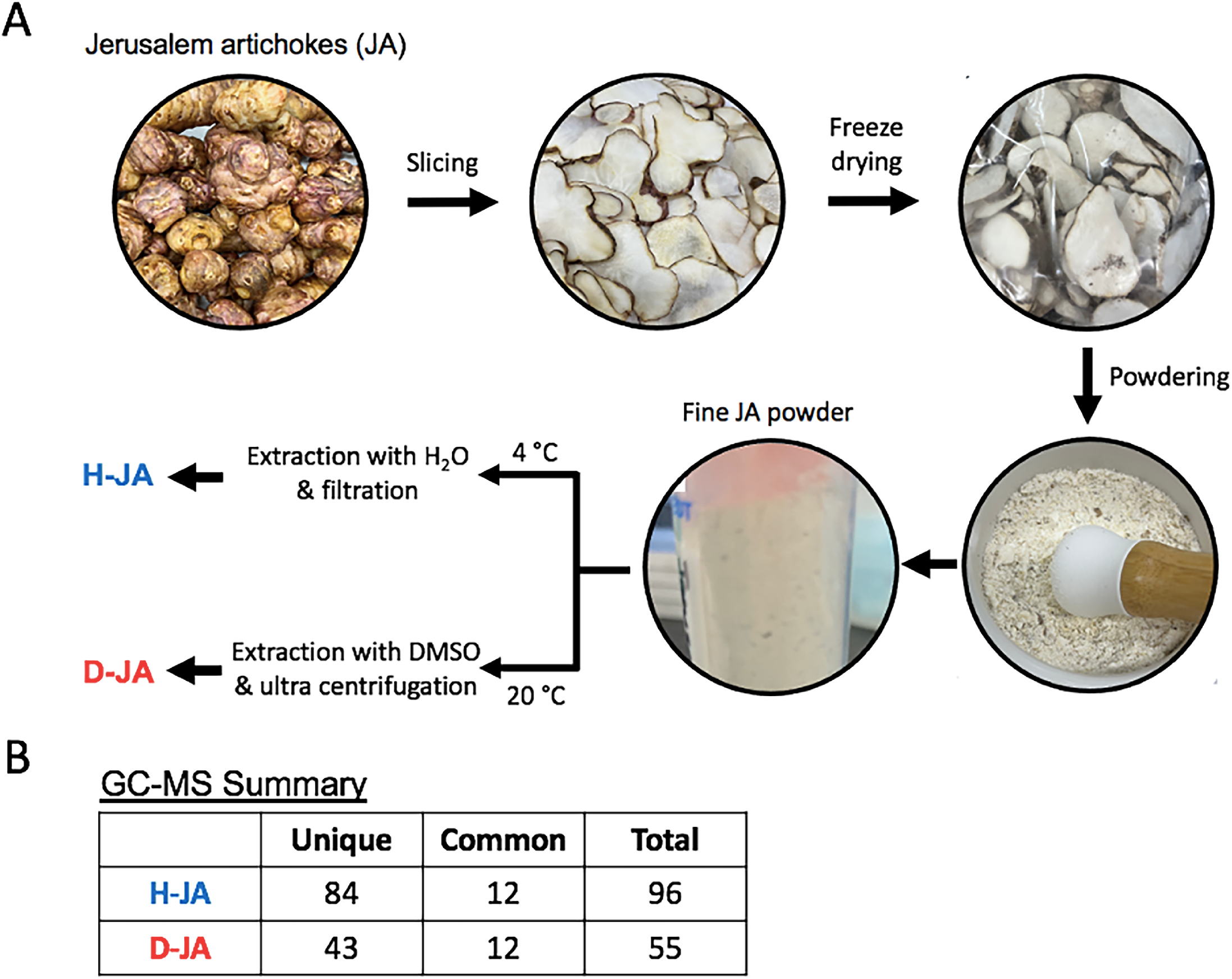
Generation of water- and DMSO-soluble JA extracts. **(A)** A schematic representation of processing JA to generate H-JA and D-JA. **(B)** Summary of the numbers of chemical compounds identified with H-JA and D-JA through GC-MS analyses.

### D-JA reduces *FASN* gene expression in SH-SY5Y cells

**T**he effects of H-JA and D-JA on *FASN* gene expression were analyzed in the SH-SY5Y human bone marrow cell line. *FASN* is expressed in a tissue-specific manner: with higher levels detected in certain tissues, including fat, brain, liver, and colon. It is also expressed in small amounts in the bone marrow, kidney, and skin^25^. H-JA and D-JA were supplemented to SH-SY5Y cells at final concentrations of 0.01% and 0.1%, with the same amounts of H_2_O and DMSO as the controls (CTRL) for 48 h. The cells were analyzed using the WST assay. The results showed that 0.1% H-JA stimulated cell growth, whereas 0.01% H-JA did not. D-JA supplementation at the indicated concentrations rarely affected cell proliferation rates compared with CTRL (**Fig. 2A**).

**Fig. 2.**
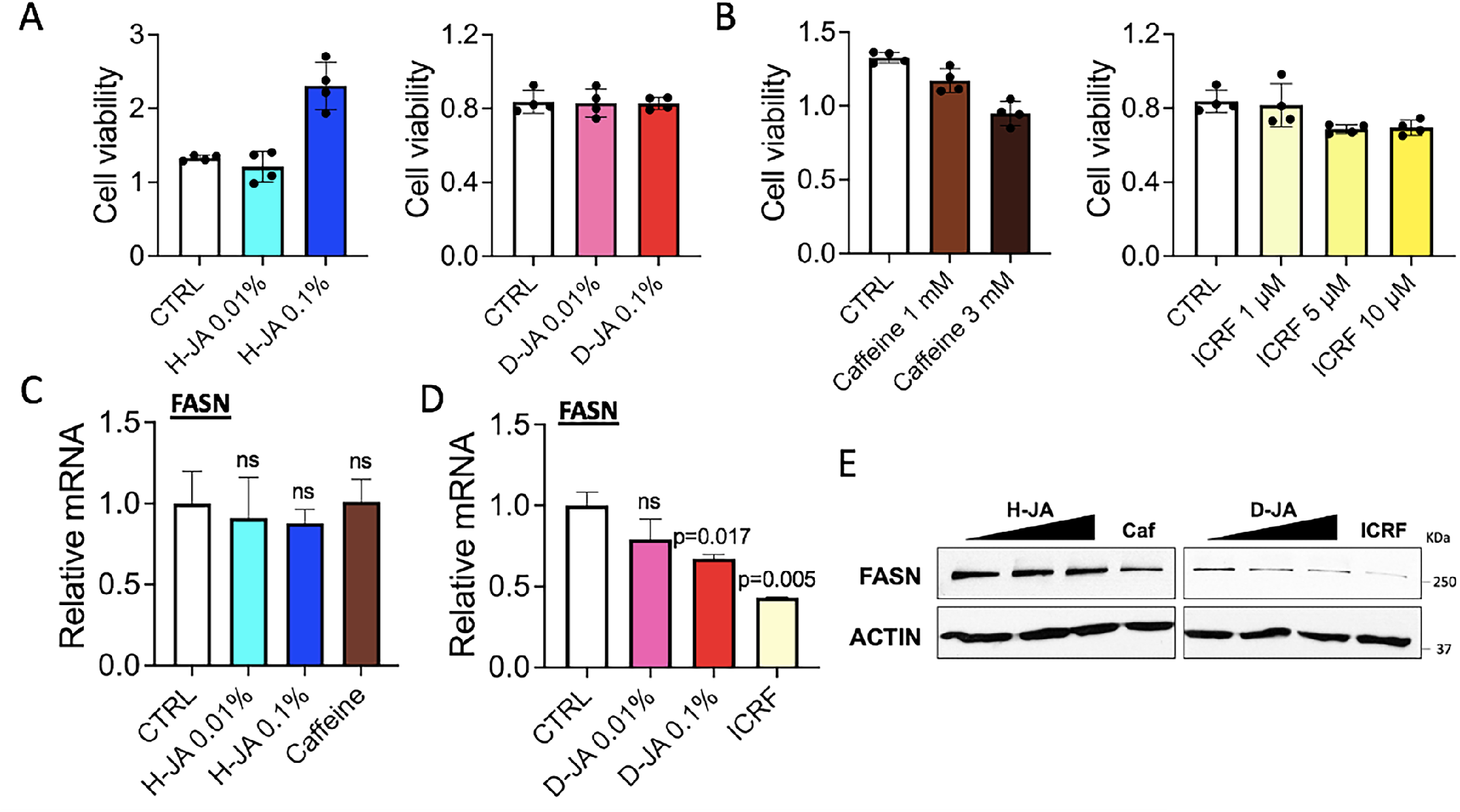
D-JA reduces *FASN* gene expression in SH-SY5Y cells. **(A)** Results of the WST cell viability assay with SH-SY5Y cells. H-JA and D-JA were supplemented to the cells for 48 h. CTRL for H-JA and D-JA, H_2_O and DMSO, respectively. Y axis, absorbance at 450 nm to measure the amount of formazan. n = 4. **(B)** Results of the WST cell viability assay with SH-SY5Y cells. Caffeine and ICRF were supplemented to the cells for 48 h. CTRL for caffeine and ICRF, H_2_O and DMSO, respectively. Y axis, absorbance at 450 nm to measure the amount of formazan. n = 4. **(C)** qRT**–**PCR results showing the effect of H-JA and caffeine on the *FASN* mRNA expression in SH-SY5Y cells. n ≥ 3. Unpaired Student’s one-tailed *t-test* was applied. ns, non-significant throughout the figures. **(D)** qRT**–**PCR results showing the effect of D-JA and ICRF on the *FASN* mRNA expression in SH-SY5Y cells. n ≥ 3. Unpaired Student’s one-tailed *t-test* was applied. **(E)** Immunoblotting results showing the effect of H-JA, D-JA, caffeine (Caf), and ICRF on FASN protein levels in SH-SY5Y cells. H-JA and D-JA were treated at final concentrations of 0, 0.01, and 0.1%. β-ACTIN was used as a loading control.

We wanted to analyze caffeine and ICRF193 (ICRF, hereafter) as positive controls for comparison with JA extracts. Caffeine plays a role in suppressing adipogenesis and *FASN* expression in fat and hepatic cells and mice^26-28^. ICRF is a catalytic inhibitor of TOP2. According to a previous study, TOP2B and DNA-PK are important regulators of *FASN* transcription^24^. Therefore, we hypothesized that inhibiting TOP2B using ICRF may interfere with *FASN* transcription. To set up the concentrations to treat the cells, the cytotoxicity of 1 and 3 mM caffeine dissolved in H_2_O and 1, 5, and 10 µM ICRF dissolved in DMSO was tested using the WST assay. The data indicated noticeable cytotoxicity with 3 mM caffeine and mild cytotoxicity with 1 mM caffeine and 5 and 10 µM ICRF (**Fig. 2B**).

Next, H-JA and D-JA along with 1 mM caffeine and 5 µM ICRF were compared for their effects on *FASN* mRNA expression in SH-SY5Y cells. JA samples and the controls were supplemented to the cells for 48 h before performing qRT**–**PCR analyses. H-JA and caffeine at the tested concentrations did not affect *FASN* transcription (**Fig. 2C**). Conversely, 0.1% D-JA significantly reduced *FASN* mRNA expression (**Fig. 2D**). As expected, ICRF potently decreased the mRNA expression of *FASN*, which confirmed the crucial role of TOP2 in *FASN* transcription (**Fig. 2D**). Under the same experimental conditions, FASN protein expression was analyzed via immunoblotting. The results showed that D-JA, as well as ICRF, but not H-JA, effectively downregulated FASN protein levels, consistent with the qRT**–**PCR results (**Figs. 2D,E**). In addition, caffeine (or Caf in immunoblotting figures) showed a mild decrease in FASN protein levels (**Fig. 2E**).

### D-JA reduces *FASN* gene expression in HCT116 cells

Next, the JA extracts were tested in the HCT116 human colon cell line. *FASN* is relatively abundantly expressed in the colon^25^, and we reasoned that the consumed JA compounds could be in direct contact with colonic epithelial cells. After 48 h, the cytotoxicity of H-JA and D-JA was evaluated using the WST assay. As shown in **Fig. 3A** and **3B**, neither H-JA nor D-JA up to a concentration of 0.2% exhibited toxicity to the cells.

**Fig. 3.**
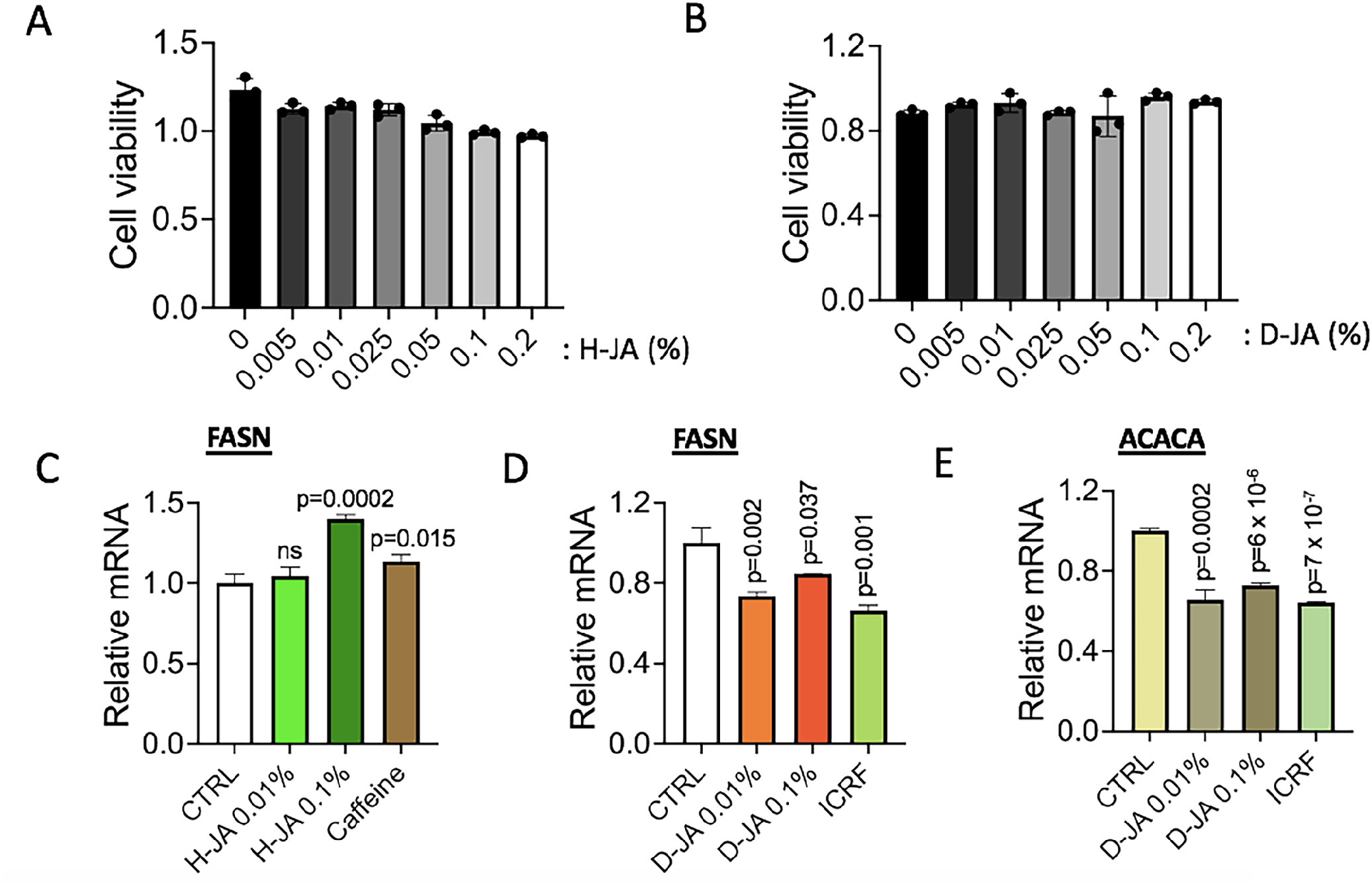
D-JA reduces *FASN* gene expression in HCT116 cells. **(A)** Results of the WST cell viability assay with HCT116 cells. H-JA was supplemented to the cells for 48 h. Y axis, absorbance at 450 nm to measure the amount of formazan. n = 3. **(B)** Results of the WST cell viability assay with HCT116 cells. D-JA was supplemented to the cells for 48 h. Y axis, absorbance at 450 nm to measure the amount of formazan. n = 3. **(C)** qRT**–**PCR results showing the effect of H-JA and caffeine on the *FASN* mRNA expression in HCT116 cells. n ≥ 3. Unpaired Student’s one-tailed *t-test* was applied. **(D)** qRT**–**PCR results showing the effect of D-JA and ICRF on the *FASN* mRNA expression in HCT116 cells. n ≥ 3. Unpaired Student’s one-tailed *t-test* was applied. **(E)** qRT**–**PCR results showing the effect of D-JA and ICRF on the *ACACA* mRNA expression in HCT116 cells. n ≥ 3. Unpaired Student’s one-tailed *t-test* was applied.

qRT**–**PCR data showed a slight increase in *FASN* mRNA expression in the presence of H-JA (0.1%, **Fig. 3C**) and caffeine. Conversely, *FASN* mRNA levels were significantly decreased by 0.01% and 0.1% D-JA and ICRF (**Fig. 3D**). The suppressive effect of D-JA was consistent with that of FASN protein expression (**Supplementary Figure 1**). The mRNA expression of *ACACA*, another key enzyme involved in fatty acid biosynthesis^29^, was measured using qRT–PCR. Interestingly, D-JA and ICRF reduced the *ACACA* mRNA levels (**Fig. 3E**), suggesting a potent effect of D-JA on suppressing both key genes involved in fatty acid biosynthesis.

### D-JA reduces *FASN* gene expression in HepG2 cells

The liver is a central organ for metabolisms in human body. Fatty acids are converted into triglycerides, which is a major form of storage fats^29^. Hepatocytes typically store small amounts of triglycerides, however, in obesity and overnutrition, hepatic fatty acid metabolism accumulates large amounts of triglycerides in hepatocytes^30^. This clinical condition is referred to as non-alcoholic fatty liver disease^31^. Given the important role of the liver in fatty acid biosynthesis, we attempted to test the effects of JA extracts on the *FASN* and *ACACA* expression in the HepG2 human hepatocyte cell line. H-JA and D-JA were added to the cells at final concentrations of 0, 0.005, 0.01, 0025, 0.05, 0.1, and 0.2%, and the WST assay was performed. At these tested concentrations, both H-JA and D-JA appeared to be innocuous (**Figs. 4A,B**). Caffeine and ICRF were used at 1 mM and 5 µM, respectively, and unlike the results in SH-SY5Y cells, they exhibited notable cytotoxicity to HepG2 cells (**Figs. 4A,B**).

**Fig. 4.**
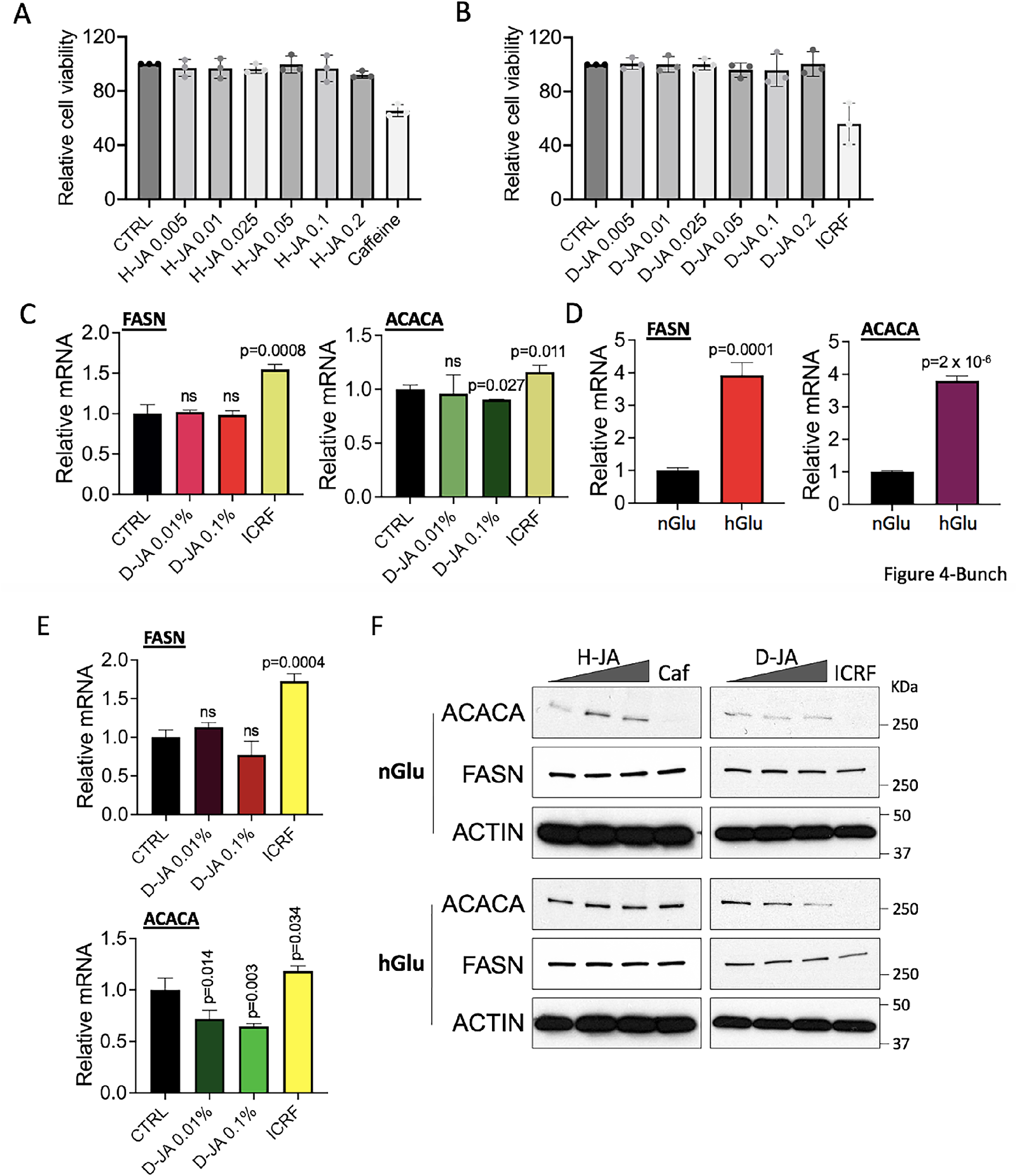
D-JA reduces *FASN* gene expression in HepG2 cells. **(A)** Results of cell viability assay using WST in HepG2 cells. H-JA was supplemented to the cells at final concentrations of 0.05, 0.01, 0.025, 0.05, 0.1, and 0.2% for 48 h. CTRL, H_2_O. Caffeine was treated at a final concentration of 1 mM. Y axis, relative absorbance (450 nm) values to CTRL as 100%. n = 3. **(B)** Results of cell viability assay using WST in HepG2 cells. D-JA was supplemented to the cells at final concentrations of 0.05, 0.01, 0.025, 0.05, 0.1, and 0.2% for 48 h. CTRL, DMSO. ICRF was treated at a final concentration of 5 µM. Y axis, relative absorbance (450 nm) values to CTRL as 100%. n = 3. **(C)** qRT**–**PCR results showing the effect of D-JA and ICRF on the mRNA expression of *FASN* and *ACACA* genes in HepG2 cells. n ≥ 3. Unpaired Student’s one-tailed *t-test* was applied. **(D)** qRT**–**PCR results showing the effect of glucose on the mRNA expression of *FASN* and *ACACA* genes under normal (nGlu, 1000 mg/L) vs high glucose (hGlu, 4500 mg/L) conditions in HepG2 cells. n ≥ 3. Unpaired Student’s one-tailed *t-test* was applied. **(E)** qRT**–**PCR results showing the effect of D-JA and ICRF on the mRNA expression of *FASN* and *ACACA* genes in HepG2 cells grown under hGlu-containing media. n ≥ 3. Unpaired Student’s one-tailed *t-test* was applied. **(F)** Immunoblotting results showing the effect of H-JA, D-JA, caffeine (Caf), and ICRF on FASN and ACACA protein levels in HepG2 cells. H-JA and D-JA were treated to final concentrations of 0, 0.01, and 0.1%. Caffeine (Caf) and ICRF were treated at final concentrations of 50 µM and 5 µM, respectively. β-ACTIN was used as a loading control.

When the cells were treated with 0.01% or 0.1% D-JA, the *FASN* mRNA level was not significantly altered, whereas ICRF at 5 µM unexpectedly increased *FASN* transcription (**Fig. 4C**). It is noted that ICRF inhibited *FASN* transcription in both SH-SY5Y and HCT116 cells (**Figs. 2D**,**E and 3D**). D-JA mildly decreased *ACACA* transcription at 0.1%, whereas ICRF increased it (**Fig. 4C**). Again, ICRF decreased the *ACACA* mRNA level in HCT116 cells (**Fig. 3E**). These data indicate that D-JA suppresses *ACACA* transcription in HepG2 and HCT116 cells, whereas ICRF has opposite effects on *FASN* and *ACACA* transcription in these cells.

In addition, we asked whether JA extracts could regulate *FASN* and *ACACA* expression under high-glucose conditions, which recapitulates overnutrition or hyperglycemia/diabetes *in vitro*. The cells were grown in media containing glucose either at a normal physiological concentration (1000 mg/L, nGlu) or a higher concentration (4500 mg/L, hGlu), and the mRNA levels of *FASN* and *ACACA* were compared. The qRT**–**PCR results showed that *FASN* and *ACACA* transcription was markedly activated under hGlu conditions (**Fig. 4D**). Similar to the results observed under the nGlu condition, D-JA did not reduce *FASN* mRNA levels, whereas ICRF increased *FASN* mRNA levels under the hGlu condition (**Fig. 4E**). Alternatively, D-JA dose-dependently decreased the mRNA levels of *ACACA* under hGlu-induced growth conditions (**Fig. 4E**). Immunoblotting confirmed that D-JA reduced ACACA protein expression in HepG2 cells under both nGlu and hGlu conditions (**Fig. 4F**). Although ICRF increased the mRNA levels of *ACACA*, it dramatically decreased the protein levels of ACACA, suggesting the formation of abnormal transcripts or translational interference by the chemical (**Figs. 4C,E,F**).

### TOP2B, a novel transcription factor of *ACACA* gene, is regulated by glucose level and D-JA

To understand the function of TOP2B in *FASN* and *ACACA* expression, we analyzed the RNA-seq data of TOP2B knockout (KO) SH-SY5Y cells (GSE142383)^32^ and evaluated the effect of TOP2B KO on the mRNA production of these genes. *FASN* was rarely affected (**Supplementary Figure 2**), whereas *ACACA* displayed a mild yet significant reduction in the absence of TOP2B (**Fig. 5A**). As a previous study reported the important role of DNA-PK in *FASN* transcription^24^, we also monitored *FASN* and *ACACA* transcription in DNA-PK–KO HCT116 cells (gifted by A. Davis Laboratory at the University of Texas Southwestern Medical Center, USA)^33^. The results illustrated that *FASN* and *ACACA* mRNA expression was decreased in DNA-PK KO cells (**Fig. 5B**). Interestingly, DNA-PK KO reduced TOP2B protein levels in immunoblotting (**Fig. 5C**). We hypothesized that decreased TOP2B activity or protein levels interfered with *FASN* and *ACACA* transcription, as shown by the results in ICRF–treated or TOP2B KO cells (**Figs. 2E, 3D,E, 4F, and 5A,B**). However, the hGlu condition dramatically reduced TOP2B protein levels and increased ACACA protein levels (**Fig. 5D**; ACACA blot is the same as one in **Fig. 4F** with different exposures). Under nGlu conditions, D-JA treatment increased TOP2B levels but decreased ACACA protein levels (**Figs. 4C,F, 5D**). These data suggest multilayer interactions and mechanisms that regulate *FASN* and *ACACA* expression. Abnormal TOP2B activities or levels may disturb the required interactions and mechanisms to up- or down-regulate *FASN* and *ACACA* expression. In addition, our data indicated that D-JA increased cellular TOP2B levels, which resulted in deregulated *ACACA* gene expression.

**Fig. 5.**
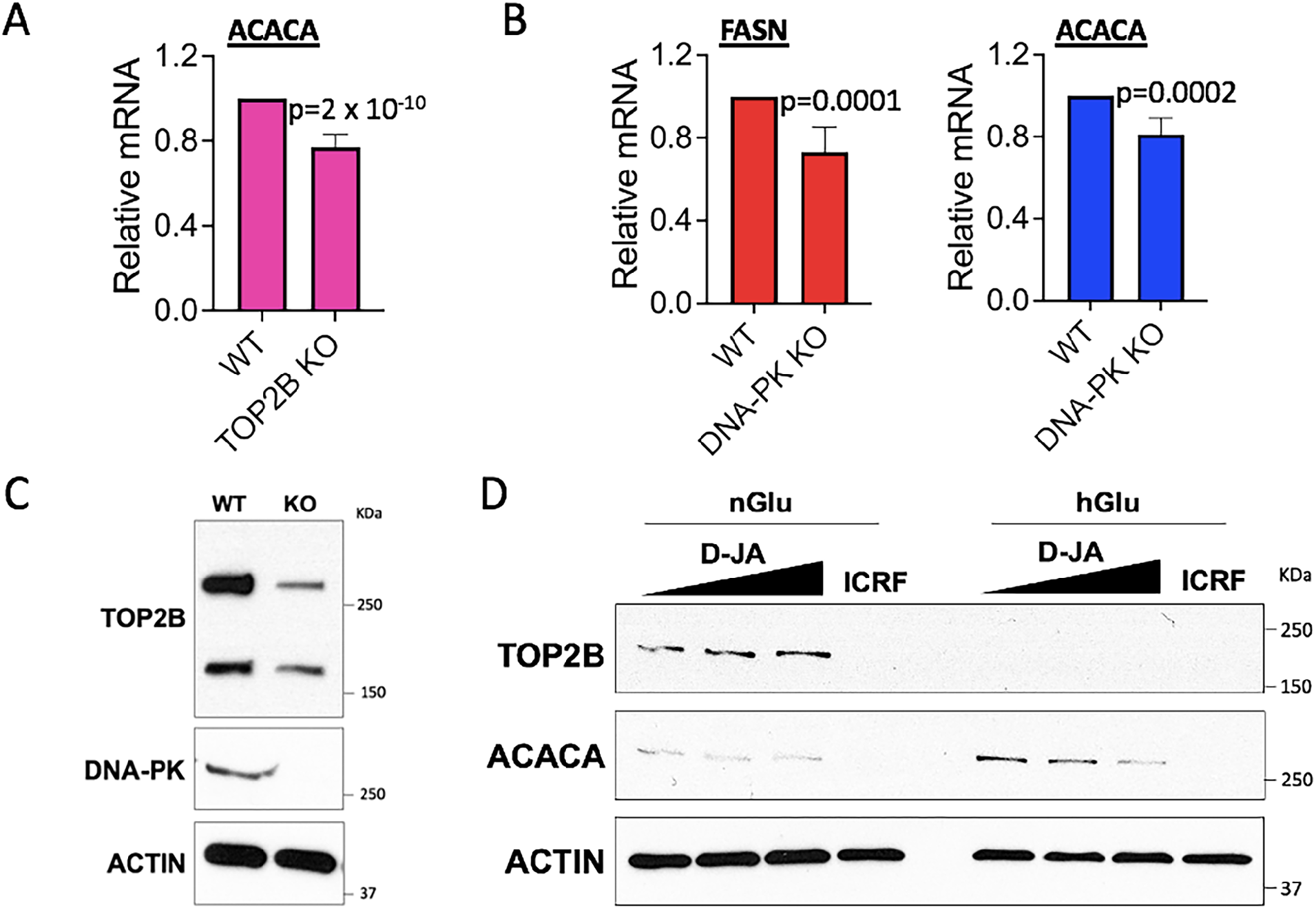
TOP2B, a novel transcription factor of *ACACA* gene, is regulated by glucose level and D-JA. **(A)** RNA-seq results for *ACACA*, comparing WT and TOP2B KO**–**SH-SY5Y cells. Unpaired Student’s one-tailed *t-test* was applied. **(B)** qRNA**–**PCR results for *FASN* and *ACACA* genes, comparing WT and DNA-PK**–**KO HCT116 cells. Unpaired Student’s one-tailed *t-test* was applied. n = 3. **(C)** Immunoblotting results showing TOP2B reduction in DNA-PK**–**KO HCT116 cells. β-ACTIN was used as a loading control. **(D)** Immunoblotting results showing the effect of high glucose (hGlu) on TOP2B protein levels in HepG2 cells. Although both hGlu and ICRF markedly reduced TOP2B protein level, hGlu stimulated *ACACA* expression, whereas ICRF under both nGlu (normal glucose) and hGlu conditions decreased it. D-JA increased TOP2B under nGlu conditions but decreased *ACACA* expression both under nGlu and hGlu conditions. β-ACTIN was used as a loading control.

## DISCUSSION

In this study, we identified a novel function of D-JA in suppressing *FASN* and *ACACA* gene expression in SH-SY5Y, HCT116, and HepG2 cells. Although both H-JA and D-JA were not cytotoxic, H-JA did not exhibit any notable, consistent effects on the expression of these genes. Note that inulin, a fructose polymer, is water-soluble. This well-known health beneficial polar compound is expected to be extracted by both H_2_O and DMSO, which suggest that the active compound(s) in D-JA that suppresses *FASN* and *ACACA* genes might be different one(s). Our GC-MS data identified common and unique compounds in H-JA and D-JA. However, the lack of established chemical libraries for JA hindered the identification of potential effector molecules. Additionally, GC analysis preferably separates volatiles and thus, most polar compounds are rarely detected. The effector molecules should be identified in consecutive studies, and the role of D-JA in suppressing *FASN* and *ACACA* expression awaits further studies including *in vivo* experiments in future.

On the other hand, crude extracts of natural products have unique health and economic benefits as they contain effectors under near-native conditions and require less processing and refinement than synthetic chemicals. Recently, we have reported the novel and potent function of *Lactiplantibacillus plantarum K8* lysates from Korean fermented cabbage called kim-chi in suppressing hypoxia-inducible gene expression^34^. More and more studies have reported medicinal and functional chemicals in foods and natural products and many of these have been developed into medicines, health supplements, and commodities^35,36^. It is not surprising that natural chemicals, just similar to synthetic chemicals, can function as environmental signals or cellular messengers that regulate gene expression^37,38^.

Although JA has been used as a remedy to lower blood sugar levels for many years, our study for the first time identified that it can suppress transcription of the *FASN* and *ACACA* genes, suggesting that JA regulates fatty acid and triglyceride biosynthesis (**Fig. 6**). The new function of JA involves TOP2B modulation. We report that TOP2B catalytic activity is required for the proper transcription of *FASN* and *ACACA* and that inhibition of TOP2B catalysis by ICRF deregulates it (**Figs. 2D,E, 3D,E, 4C,E,F**, and **5D**). We found that cellular TOP2B protein level has complex impacts on *FASN* and *ACACA* expression. Although TOP2B KO tends to decrease the transcription activities at these genes, high-glucose concentrations, which reduces TOP2B protein levels, increase both *FASN* and *ACACA* expression (**Fig. 5**). Likewise, D-JA increases TOP2B protein levels and decreases *ACACA* expression in HepG2 cells (**Figs. 4D–F, 5D, 6**). It should be validated whether TOP2B-gene (DNA) interaction is interfered by D-JA, whether D-JA inhibits TOP2B catalytic activity, or both. In addition, it seems reasonable to consider that genomic topological integrity is guarded by topoisomerases; however, their excess or insufficient activities might burden the DNA, thereby inhibiting DNA metabolism, including transcription.

**Fig. 6.**
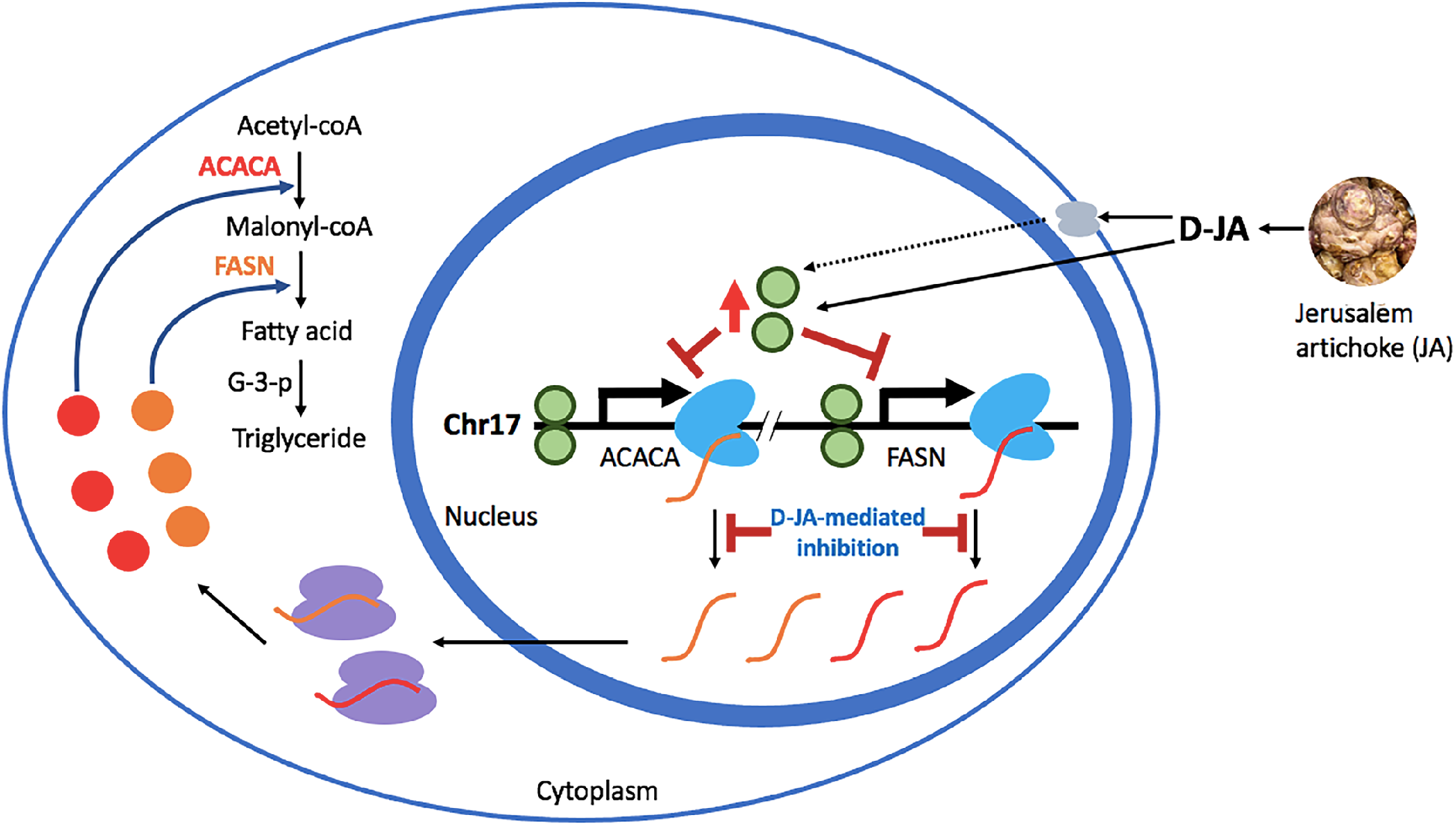
A model of D-JA-mediated *FASN* and *ACACA* gene regulation. Both *ACACA* and *FASN* genes, key enzymes involved in fatty acid biosynthesis, are located in chromosome 17 (Chr17). Transcribed mRNAs (red curved lines for *ACACA*; orange curved lines for *FASN*) are exported to cytoplasm and translated into proteins (red circles for ACACA protein; orange circles for FASN protein) by ribosomes (purple circular dimers). ACACA converts acetyl-coA released from mitochondria into malonyl-coA and FASN synthesizes fatty acid. Glycerol-3-phosphate (G-3-P) and fatty acids are covalently complexed through multiple steps to generate triglyceride, a major form of storage fats in humans^29^. Undefined compounds in JA-derived D-JA may directly enter the cell and nucleus (marked with a solid black line) to affect gene expression or/and may interact with membrane proteins (gray ovals) to propagate the signals inside the cell (marked with a dotted black line). D-JA deregulates cellular TOP2B protein (green circles) levels and suppresses the transcription of *FASN* and *ACACA* genes. Sky blue objects, RNA polymerase II transcribing the genes.

The novel function of D-JA in reducing *FASN* and *ACACA* expression identified in this study sheds light on the development of a safe and effective drug or health supplement for preventing and alleviating metabolic syndrome, including obesity, hypertension, and diabetes. In particular, along with its established function in reducing blood glucose levels^15,17,18,39^, the potential role of D-JA in suppressing fatty acid biosynthesis is promising for effectively treating diabetes caused by insulin resistance, which results from excess fats in the blood and interstitial fluid.

## Acknowledgments

We thank N. Dinh, J. Lee, S. Ju, B. Kang, J. Jeong and the current members of the Bunch laboratory members at Kyungpook National University (KNU) for their technical assistance. We appreciate A. J. Davis and H. Lu at the University of Texas Southwestern Medical Center for sharing their DNA-PK KO HCT116 cells. H.B. thanks J. Christ and John and D. Y. Bunch for their loving encouragement and support throughout the course of this work.

## Financial Supports

This research was supported by grants from the National Research Foundation (NRF) of the Republic of Korea (2022R1A21003569) to H.B.

## Author Contributions

HB generated JA samples and carried out cell culture, cytotoxicity analysis, qRT**–**PCR, and immunoblotting. YY and HB performed and analyzed GC-MS. SC and HB revised the manuscript. HB created the hypothesis, designed the experiments, analyzed and curated the data, and wrote the manuscript.

## Data and Materials Availability

All data are available in the manuscript or as supplementary information.

---

**Supplementary Table 1.**
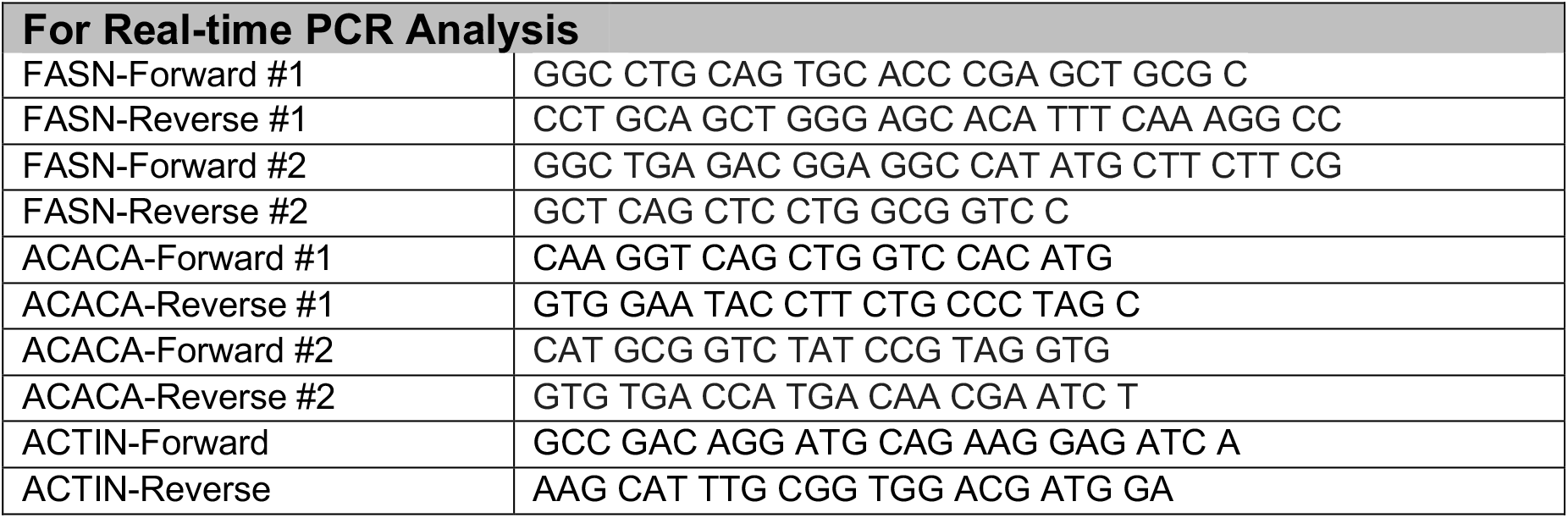
The sequences of primers used in this study.

